# Dysregulation of temporal dynamics of synchronous neural activity in adolescents on autism spectrum

**DOI:** 10.1101/813519

**Authors:** Evie A. Malaia, Sungwoo Ahn, Leonid L Rubchinsky

**Affiliations:** Department of Communicative Disorders, University of Alabama, Tuscaloosa, AL; Department of Mathematics, East Carolina University, Greenville, NC; Department of Mathematical Sciences, Indiana University Purdue University Indianapolis, Indianapolis, IN; Stark Neurosciences Research Institute, Indiana University School of Medicine, Indianapolis, IN

**Author notes:** Correspondence: E.A. Malaia.

**Keywords:** Autism Spectrum Disorder, resting state, neural oscillations, neural synchronization, functional connectivity, phase synchrony, developmental cognitive neuroscience

## Abstract

Autism spectrum disorder is increasingly understood to be based on atypical signal transfer among multiple interconnected networks in the brain. Relative temporal patterns of neural activity have been shown to underlie both the altered neurophysiology and the altered behaviors in a variety of neurogenic disorders. We assessed brain network dynamics variability in Autism Spectrum Disorders (ASD) using measures of synchronization (phase-locking) strength, and timing of synchronization and desynchronization of neural activity (desynchronization ratio) across frequency bands of resting state EEG. Our analysis indicated that fronto-parietal synchronization is higher in ASD, but with more short periods of desynchronization. It also indicates that the relationship between the properties of neural synchronization and behavior is different in ASD and typically developing populations. Recent theoretical studies suggest that neural networks with high desynchronization ratio have increased sensitivity to inputs. Our results point to the potential significance of this phenomenon to autistic brain. This sensitivity may disrupt production of an appropriate neural and behavioral responses to external stimuli. Cognitive processes dependent on integration of activity from multiple networks may be, as a result, particularly vulnerable to disruption.

**Lay Summary:** Parts of the brain can work together by synchronizing activity of the neurons. We recorded electrical activity of the brain in adolescents with autism spectrum disorder, and then compared the recording to that of their peers without the diagnosis. We found that in participants with autism, there were a lot of very short time periods of non-synchronized activity between frontal and parietal parts of the brain. Mathematical models show that the brain system with this kind of activity is very sensitive to external events.

## Introduction

Human behavior in health and disease is undergirded by temporal synchronization of distributed networks, which allows for information processing (e.g., Buzsaki, 2006; Harris & Gordon, 2015). The difficulty in understanding of the temporal regulation of brain networks is partially due to the disparity of temporal scales for different types of neural data (e.g. between millisecond scale of brain-activity-driven EEG and second-to-minute scale of blood-flow-based fMRI).

Experimental data from different modalities, such as EEG, MEG, and fMRI, indicate that better understanding of temporal dynamics of brain activity may help elucidate mechanisms of brain network organization in ASD population. During rest, ASD participants show increased coherence (or, static overconnectivity) in long-range brain networks (Buckley et al., 2015). However, the increase in coherence coincides with longer dwell times in a globally-disconnected state (Rashid et al., 2018), and increased variability in connectivity over time (Mash et al., 2019). Analyses of EEG and MEG recordings, which provide higher temporal resolution, indicate long-range functional underconnectivity (O’Reilly et al., 2017), and decreased synchrony in short- and medium-range connections in ASD participants (Schwartz et al., 2017), particularly in higher frequency bands. EEG data also point to changes in brain network organization developmentally: e.g. between 3 and 11 years of age decreased synchronization among brain regions persists, but in addition to it, increases in within-regional synchronization are observed (Han et al., 2017; Kang et al., 2019). The power of the signal in ASD participants appears to be enhanced in both low- and high-frequency bands (what Wang et al., 2013 called a U-shaped function), particularly in the left hemisphere. The mixed conclusions provided by the earlier results highlight the necessity of applying dynamic approaches to understand the nuanced transient patterns of oscillatory brain activity in ASD.

Short-term patterns of brain activity can contribute to long-term connection strengthening, and, as a result, changing anatomical organization of brain networks (Chu et al., 2012). The effects of temporal patterns of synchronization vs. averaged temporal dynamics on long-term brain network organization have not yet been investigated.

Research in neurobiological bases of ASD has indicated that the severity of ASD-associated behaviors is correlated with large-scale connectivity abnormalities in brain networks (Di Martino et al., 2009). Accurate understanding of fine-grained temporal organization is needed to resolve conflicting findings of anatomical hypo-and hyper-connectivity in ASD participants. In the present analysis, we investigate resting state activity in adolescents with ASD using two different measures of EEG synchronization. One is the average synchronization strength between two recorded signals. The other takes into account the temporal structure of synchronization, comparing the relative duration of synchronized and non-synchronized time intervals. The two measures are conceptually different and can vary independently of each other: it is possible to have the same synchronization strength either with few long desynchronizations or with many short desynchronizations; these two opposites would lead to different behavior.

The present analysis assesses the relationship between temporal parameters of the resting state EEG, comparing adolescents on autism scale, and their neurotypical peers. The resting state activity of the brain reflects typical patterns of neural organization by activating networks that are salient for everyday existence. Characterizing temporal variability of synchronizations across brain networks provides a more faithful measure of neural activity than average synchrony strength as the latter provides only an averaged measure missing potential temporal variability. Temporal variability in synchronization can help us infer underlying changes in circuit activity (Park et al., 2010, Ahn et al., 2014). The purpose of this study is to investigate brain network dynamics in ASD, using measures of synchronization (γ, phase-locking strength averaged over time) and desynchronization (DR, desynchronization ratio) of neural activity. In particular, we are focusing on the dynamic variability of neural activity. A smaller DR means longer periods of desynchronization between the source electrodes. The question of difference between the strength of neural synchronization (averaged over time), and the temporal patterning of this synchronization is key to understanding information transfer in neural circuits (Lowet et al., 2016).

## Methods

### Participants

Fourteen adolescents (2 F) with diagnoses of Asperger syndrome or high-functioning ASD were recruited from a special-need school. 10 typically developing (TD) age-matched participants (4 F) were recruited from the schools of Arlington School District, TX (see Table 1 for demographics; although every effort was made to match the ages of participants in the two groups, we do not assume that developmental trajectories prior to the experiment were similar between groups). The study was approved and conducted in accordance with the ethical standards of the University of Texas at Arlington IRB, and of the Declaration of Helsinki. All parents and children provided their written informed consent/assent. The Test of Pragmatic Language-2 (TOPL-2, Phelps-Terasaki & Phelps-Gunn, 2007) was administered to each participant (TOPL evaluates social communication in context, indicating how well participants can choose appropriate content, express feelings, and handle other aspects of pragmatic language; average score on the test is 10; scores between 7 and 10 are considered low average, scores in the range of 10 to13 – high average). Students in ASD group were relatively high-functioning adolescents who, based on IQ >80, qualified for admission to special education program for those on autism spectrum, which required ADOS or ADI-R score as assessed by a certified clinically reliable health professional.

**Table 1.**
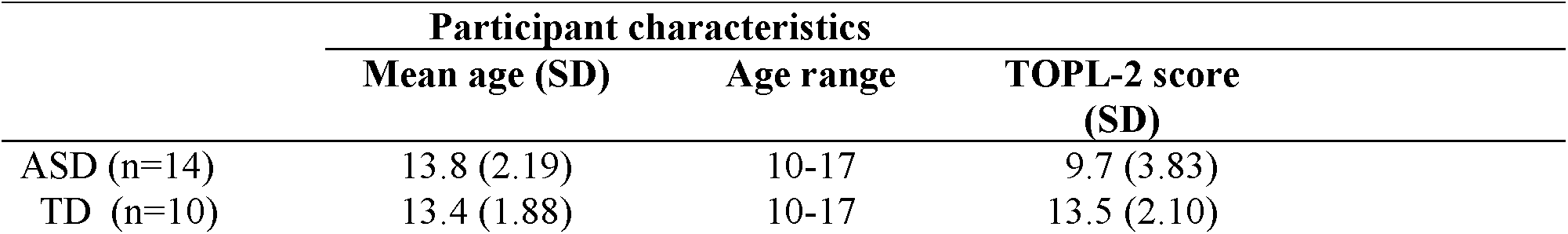
Mean age and TOPL-2 scores as a function of diagnosis. TOPL-2 scores differed significantly between groups [t(23) = −3.148, p =0.002].

### Procedure

During the EEG session, the participants were seated comfortably in a sound-attenuating booth and were asked to relax with their eyes closed. Scalp EEG was recorded from Ag/AgCl electrodes mounted in an electrode cap (Wavegard, ANT Inc.) and positioned according to the 10–20 system. Electrode impedances were maintained below 10KOhm, consistent with manufacturer’s instructions. The average recordings lasted for 97±22 sec (ASD) and 99±33 sec (TD). Synchronization analysis was performed on bilateral anterior and parietal electrodes (F3/4, P3/4), as several investigations implicated frontal and parietal networks in resting-state abnormalities in ASD (Ghanbari et al., 2015; Kitzbichler et al., 2015; Shou et al. 2018). Selected electrodes preserved hemispheric asymmetry and were substantially remote from each other to minimize a cross-talk between signals needed to analyze the temporal patterns of synchronization.

### Data analysis

EEG data were sampled at 512 Hz, band pass filtered (using zero-phase filtering to avoid phase distortions) at 0.1–512 Hz, and referenced to the average of mastoids. The synchronization analysis methods used here were described previously in detail (Park et al., 2010; Ahn & Rubchinsky, 2013; Ahn et al., 2014). Briefly, signals were Kaiser windowed and digitally filtered using a finite impulse response filter in four bands: theta (4-7 Hz), alpha (8-12 Hz), beta (13-30 Hz), and low gamma (31-59 Hz). Hilbert transform was used to reconstruct phases of oscillations. To detect oscillatory episodes we used the signal-to-noise ratio criterion as described in (Park et al., 2010). The reconstructed phases were used to estimate the phase-locking strength computed in a conventional way:

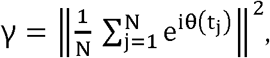

where θ is the difference of the phases of oscillations. This index varies from 0 to 1 (perfect phase synchronization) (Pikovsky et al., 2001; Hurtado et al., 2004).

The phase-locking index γ characterizes the synchronization strength averaged over the analysis window. If the oscillations are synchronized on the average, then it is possible to check whether oscillations are in the synchronized state at a specific cycle of oscillations. The recent developments in this area allowed researchers to describe the temporal patterning of neural synchronization on very short temporal scales (Park et al., 2010; Ahn et al., 2011, 2014; Ahn & Rubchinsky, 2013). Following these studies, we characterize the temporal patterning of synchronization using the distribution of desynchronization durations. This approach extracts intervals during which the phase difference is close to the preferred value, as well as intervals during which the phase difference substantially deviates from the preferred value (desynchronizations). We briefly outline this analysis and refer to the aforementioned studies for the detailed description. Whenever the phase of one signal crosses zero level from negative to positive values, we record the phase of the other signal, generating a set of consecutive values {*ϕ*_*i*_}, *i*=1,…, *N*. These *ϕ*_*i*_ represent the phase difference between two signals (Fig. 1C). After determining the most frequent value of *ϕ*_*i*_, all the phases are shifted accordingly (for different episodes under consideration) so that averaging across different episodes (with potentially different phase-shifts) is possible. Thus, this approach is not concerned with the value of the phase shift between signals, but rather with the maintenance of the constant phase-shift (phase-locking). Temporal dynamics are considered to be desynchronized if the phase difference deviates from the preferred phase difference by more than π/2 as in the earlier studies. The duration of the desynchronized episodes is measured in cycles of the oscillations. For example, if the phase difference deviates from the preferred phase difference by more than π/2 once, then the duration of the desynchronized episodes is one (Fig. 1C at *ϕ*_6_). If it deviates twice, then the duration is two, etc. This approach considers the maintenance of the phase difference in time and distinguishes between many short desynchronizations, few long desynchronizations, and possibilities in between even if they yield the same average synchrony strength.

**Figure 1.**
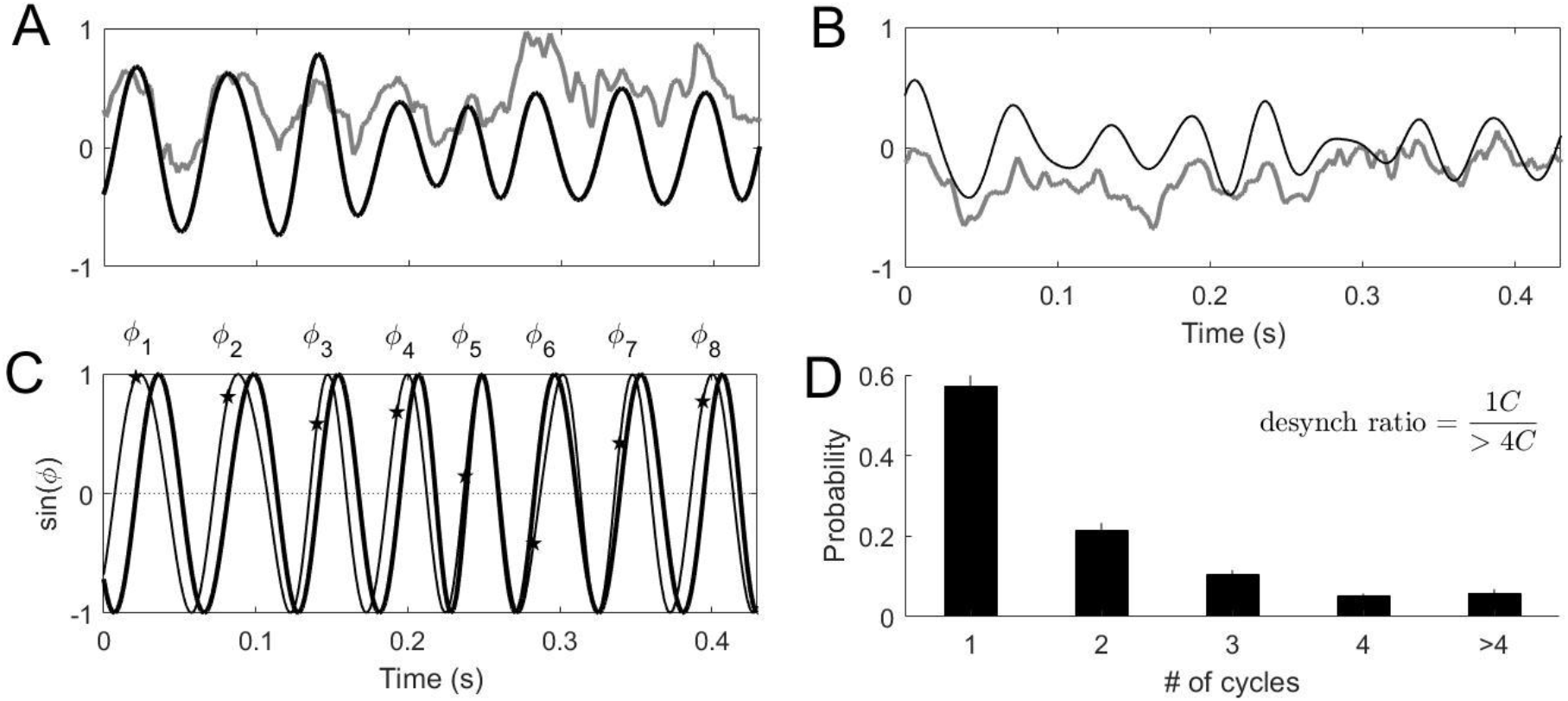
(A) and (B) are examples of the raw (gray line) and filtered (black line) EEG signals recorded from F3 and F4 electrodes respectively. (C) The sines of the phases of both filtered signals. The symbols *ϕ*_*i*_ represent the values of the phase of one signal, when the phase of the other crosses the zero from negative to positive value. The desynchronization (relative to the most common phase lag) happens around *ϕ*_6_. (D) The distributions of desynchronization durations for F3 and F4 for all patients measured in the number of cycles of oscillations (mean ±SEM was plotted).

Following Ahn et al. (2014, 2018) we evaluate DR: the ratio of the relative frequencies of the desynchronizations lasting for one cycle to longer than 4 cycles of oscillations (Fig. 1D). A smaller value of DR identifies a bias toward longer desynchronizations while a larger value of DR identifies a bias toward short desynchronizations. Importantly, the average synchronization strength can vary independently of DR (Ahn et al., 2014; Ahn & Rubchinsky, 2017): γ characterizes the average synchrony while DR describes its temporal variation. Measuring the durations of desynchronizations in cycles of oscillations (instead of actual time unit) allows for comparison of temporal patterns of synchronization between different brain rhythms.

We would like to note that DR can be considered as a measure of dynamic variability of neural activity (specifically focused on the synchronized dynamics). Other measures of dynamics variability of neural activity (methods based on the different types of entropy of the time-series) have been applied to EEG signals yielding differences between ASD and TD subjects (Kang et al., 2019). In that respect, DR is focused on the dynamical aspects of variability of specific (oscillatory) mechanisms of neural activity, not just general differences in complexity.

Pairwise γ and DR were computed between anterior (F3, F4) and parietal (P3, P4) sites. The values of γ and DR were averaged over 30 sec non-overlapping windows. All comparisons were first subjected to a mixed design analysis of variance (ANOVA) followed by two-sample two-sided t-test with significance α=0.0125 (Bonferroni correction).

We further computed Pearson correlations between TOPL-2 scores and the synchrony measures (γ and DR) of individual participants for ASD and TD groups. Fisher’s r to z transformation was applied after conducting Pearson correlation analysis between synchrony measures and TOPL-2 scores to access whether the correlations between groups differed significantly.

## Results

EEG data of ASD participants manifested altered coordination of activity in frontal and parietal regions in multiple frequency bands; moreover, temporal dynamics of this coordination was also altered (Figure 2). To investigate the difference of synchronization strength between ASD and TD, we performed a mixed design ANOVA (between-subject factor as group, and within-subject factors as frequency and location). There were significant main effects of group [F(1, 66)=7.85, p=6.66e−3], frequency [F(3, 198)=1.99e+2, p<1.0e−16], and location [F(3, 198)=6.58e+1, p<1.0e−16]. There were also significant interactions in frequency*location [F(9, 594)=3.36e+1, p<1.0e−16] and group*frequency*location [F(9, 594)=2.90, p=2.28e−3], but no interactions in group*frequency [F(3, 198)=1.77e−1, p>0.05] and group*location [F(3, 198)=5.12e−1, p>0.05]. We performed the further analyses at each frequency band and location to compare the difference of synchronization strengths between groups. Synchronization strength in fronto-parietal sites was altered in ASD in several locations at alpha band [P3-P4, t(66)=3.45, p=9.87e−4], beta band [F3-P3, t(66)=4.57, p=2.22e−5; F4-P4, t(66)=3.20, p=2.08e−3], low gamma band [F3-F4, t(66)=2.98, p=3.99e−3] while other electrode pairs did not exhibit significant difference. We observed that the synchrony strength of F3-P3 in ASD at theta band was marginally different [t(66)=2.15, p=3.55e−2] compared to TD participants. In general, this alteration was in the direction of higher synchrony in ASD compared to TD participants.

**Figure 2.**
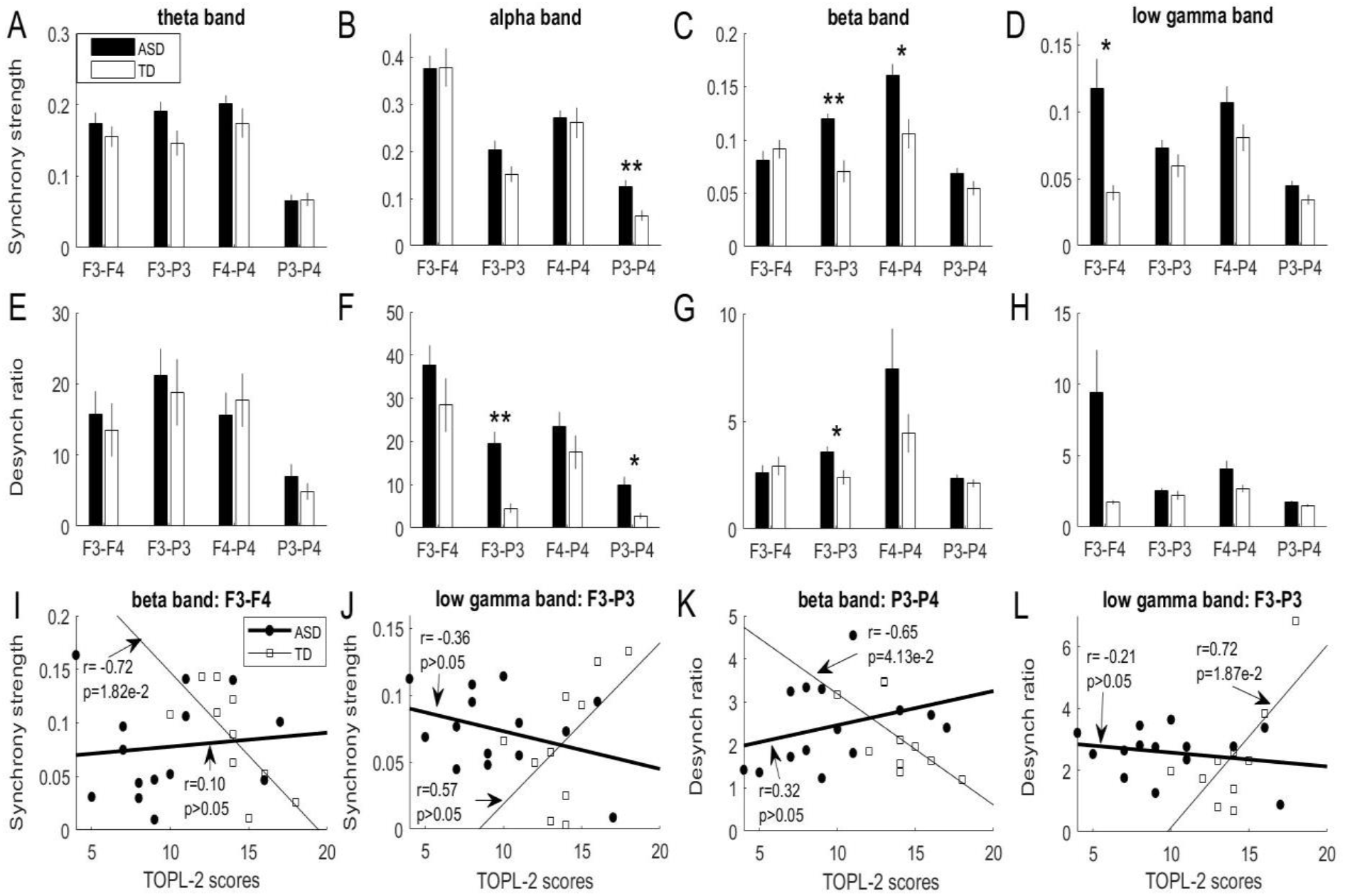
Synchronization strength γ in upper panels (A-D) and desynchronization ratio DR in middle panels (E-H) calculated pairwise across F3, F4, P3, and P4 electrodes and frequency bands in resting state EEG. Mean±SEM was plotted. Asterisks represent paired two-sided t-test (*p<0.0125; **p<0.00125). Lower panels (I-L) calculated Pearson correlations between synchronization measures (γ and desynchronization ratio) and TOPL-2 scores. (I-L) demonstrated that the correlations between synchrony measures and TOPL-2 scores at the given frequency and location significantly differed between TD and ASD groups.

Temporal patterning of synchronization of neural activity was found to be altered in ASD too. To investigate the difference of DRs between ASD and TD, a mixed design ANOVA was performed. There were significant main effects of group [F(1, 66)=8.48, p=4.90e−3], frequency [F(3, 198)=6.19e+1, p<1.0e−16], and location [F(3, 198)=2.38e+1, p=3.37e−13]. There were also significant interactions in group*frequency [F(3, 198)=4.28, p=5.94e−3], frequency*location [F(9, 594)=1.22e+1, p<1.0e−16], but no interactions in group*location [F(3, 198)=6.76e−1, p>0.05] and group*frequency*location [F(9, 594)=7.82e−1, p>0.05]. We performed the further analyses at each frequency band and location to compare the difference of DRs between groups. In particular, there was a trend for more numerous short desynchronizations in ASD subjects as detected by DR at alpha band [F3-P3, t(66)=4.39, p=4.14e−5; P3-P4, t(66)=2.89, p=5.18e−3] and beta band [F3-P3, t(66)=2.97, p=4.18e-3] while other electrode pairs did not exhibit significant difference. We also observed that DR of F3-F4 at low gamma band in ASD subjects was marginally different [t(66)=2.34, p=2.88e-2] compared to TD participants. This alteration was in the direction of more numerous short desynchronizations in ASD subjects.

To compare the difference of correlations between TOPL-2 scores and the synchrony measures (γ and DR) for ASD and TD groups, we performed Fisher’s r to z transformation (see *Data analysis* for detail). Difference of correlations analysis indicated that sign of correlation between posterior interhemispheric (P3-P4) DR in the beta range and TOPL-2 scores significantly differed between TD and ASD groups [significant negative correlation for TD, non-significant positive correlation for ASD, z-score=2.30, p=2.17e−2]. As illustrated by Figure 2, in the low gamma range, the sign of correlation between left hemisphere (F3-P3) DR and TOPL-2 scores also significantly differed between groups [significant positive correlation for TD, non-significant negative correlation for ASD, z-score=2.33, p=1.98e−2]. Correlation analysis between synchrony strength in the frontal interhemispheric (F3-F4) and TOPL-2 scores also revealed the difference between groups at beta range [significant negative correlation for TD, non-significant positive correlation for ASD, z-score=2.11, p=3.52e−2]. Finally, the type of correlation between the left hemisphere (F3-P3) synchrony strength at the low gamma band and TOPL-2 scores differed between groups [marginal positive correlation for TD, non-significant negative correlation for ASD, z-score=2.11, p=3.52e−2]. While the difference in how synchronization metrics and TOPL-2 are correlated does not exhibit very large statistical significance, it points to intriguing differences in several brain networks and spectral bands. These results in Figure 2 indicate how the relationship between properties of neural synchronization and behavior may differ in TD and ASD.

## Discussion

Understanding of the neurobiological processes that underlie brain activity abnormalities in ASD is critical for diagnostics and development of targeted interventions. In ASD adolescents, stable cross-frequency networks during resting state have been shown to endure for a longer period of time in comparison with the neurotypical peers (Malaia et al., 2016). The present analysis extends the findings to the temporal aspect of synchronization: we demonstrate that average fronto-parietal synchronization is higher in ASD, but with more short periods of desynchronization. Research on neuroanatomical structure changes in ASD has previously connected the difficulty of modulating executive control in high-functioning adolescents on autism spectrum to failure of fronto-parietal regions to function as integrative hubs within the brain’s network architecture (Lynch et al., 2017). Although the relationships between behavior and oscillatory activity were not tested directly in the present study, some relationships might be hypothesized from prior work, where decreased theta and alpha coherence were shown to lead to impairment in working memory and between-network binding (particularly as related to executive processing, inhibition, and conscious attention), while beta frequency synchrony has been related to successful higher-order cognitive processing (cf. Schwartz et al., 2017). Additionally, atypical pattern of synchronization in the left hemisphere might be related to left-lateralized microstructural abnormalities in ASD (Peterson et al., 2015)

Our analysis of fronto-parietal temporal dysregulation suggests a possible underlying mechanism whereby normal functional organization of brain networks in ASD fails to emerge. Mathematical modeling (Ahn & Rubchinsky, 2017) suggests that networks with high DR have increased sensitivity to inputs; this sensitivity may disrupt production of adequate neural and behavioral responses to external stimuli. Cognitive processes dependent on integration of activity from multiple networks may be, as a result, particularly vulnerable to disruption. With regard to the function of connectivity between frontal and parietal regions, Velazquez et al. (2009) note that disruption in phase synchronization between frontal and parietal regions in ASD participants is correlated with impaired executive function in tasks. Murias et al. (2007) also noted reduced coherence between frontal and other regions in the alpha band of resting state EEG in ASD participants, while Coben et al. (2008) indicated reduced long-range within-hemispheric coherence in beta, theta, and delta ranges.

The situation observed here (some spectral bands exhibit the difference between TD and ASD in synchrony strength, some in synchrony patterning, and some in both) should not be viewed as an anomaly (cf. Ahn et al., 2018), as alterations of synchrony patterning and synchrony strength characterize different aspects of synchronized activity and do not necessarily follow each other. Desynchronization ratio and synchronization strength characterize different aspects of neural synchrony. Thus, the results of comparison of TD and ASD synchrony indicate multiple functional differences in brain networks of TD and ASD subjects. Computational modeling suggests that higher DR leads to higher sensitivity to inputs. In a model of small network of coupled neurons, which are subjected to common synaptic input, a higher DR (even with the same average synchrony strength) leads to situation where a weaker synaptic input is needed to reach any (pre-selected) level of synchrony. Our understanding of these modeling results is that a higher DR means the network is switching between synchronized and desynchronized states more frequently (this parameter is not necessarily related to the overall synchrony strength, as the durations of synch and desynch intervals may be different). Using the language of dynamical systems theory, the system with a higher DR brings itself to the vicinity of synchronous state on its own (although synchronous state is unstable, and the system cannot stay there for a long time). Thus, in a system with high DR, synchrony does not need to be created *de novo* in the phase space, but rather it needs to be stabilized for some time, which is possible to do with weak input – thus there is more sensitivity in the system.

Finally, we note that our results showed that TD group had relatively lower synchrony strengths and lower DRs with stronger correlation with TOPL-2 scores as compared to ASD group. On the other hand, the correlations between neurophysiology and behavior in ASD subjects were weak and moved toward the opposite directions of those in TD subjects. This may suggest that multiple mechanisms may be involved in neural processing in ASD so that the same change in synchrony strength or patterning may lead to different behavioral changes in ASD and control participants. For example, greater beta synchrony strength across frontal regions seemed detrimental to language scores in TD participants, but not ASD participants. The stronger correlation between the synchrony measures and participant’s language scores in TD group may suggests that regulation of the neural oscillations in the normal brain may be effectively translated to the behavioral domain. However, this is not the case for the abnormally elevated and abnormally patterned synchronization in ASD. These findings call for a deeper future exploration of functional significance of oscillatory patterns in different clinical populations

The relatively small number of participants and their relatively high functioning status make for a limiting factor, however, they also mean that the differences between the groups found here are quite likely to be more emphasized in larger populations with large functioning differences. Yet, the small number of participants coupled with relatively large number of statistical tests indicate that conclusions of this brief report are hypotheses, which should eventually be explored in more detail. Another limitation was lack of individual information on psychiatric comorbidities, medication, and IQ.

This study is the first, to our knowledge, to show that temporal aspects of dynamics of neural synchronization on very short time-scales may be essential for understanding of organization of functional neural networks in ASD. Not only the average synchrony strength, but the way how the synchronization is distributed in time is different between TD and ASD in a specific way to brain areas and spectral bands. These findings also offer a novel perspective on neural information processing control in atypical neural systems, suggesting future directions of research into information processing across not only spatial but also temporal scales in ASD. Specifically, group differences in alpha and theta frequency bands may imply possible relationships with behavioral features of ASDs, such as differences in higher-order cognitive processing, including language. Exploring measures of synchrony that fluctuate over time can help determine how these measures relate to behavioral differences.

## Acknowledgements

Supported by NSF DMS1813819 (LLR) and NSF 1734938 (EAM).

